# NeoTCR: an immunoinformatic database of experimentally-supported functional neoantigen-specific TCR sequences

**DOI:** 10.1101/2023.02.13.528383

**Authors:** Weijun Zhou, Wenting Xiang, Jinyi Yu, Zhihan Ruan, Yichen Pan, Kankan Wang, Jian Liu

**Author notes:** These authors contributed equally to this work (Weijun Zhou, Wenting Xiang). Correspondence (Kankan Wang), (Jian Liu).

## Abstract

Neoantigen-based immunotherapy has demonstrated of long-lasting antitumor activity. Recognition of neoantigens by T cell receptors (TCRs) is considered a trigger for antitumor responses. Due to the overwhelming number of TCR repertoires in the human genome, it is challenging to computationally pinpoint neoantigen-specific TCRs. Recent studies have identified a number of functional neoantigen-specific TCRs, but the corresponding information is scattered across published literature and is difficult to retrieve. To improve access to these data, we developed the NeoTCR, an immunoinformatic database containing a unified description of publicly available neoantigen-specific TCR sequences, as well as relevant information on targeted neoantigens, from experimentally supported studies across 18 cancer subtypes. A user-friendly web interface allows interactive browsing and running of complex database queries based on numerous criteria. To facilitate rapid identification of neoantigen-specific TCRs from raw sequencing data, NeoTCR offers a one-stop analysis for annotation and visualization of TCR clonotypes, discovery of existing neoantigen-specific TCRs, and exclusion of bystander viral-associated TCRs. NeoTCR will serve as a valuable platform to study the biological functions of neoantigen-associated T-cells in anti-tumor immunity to better apply neoantigen-specific TCRs in clinics. NeoTCR is available at http://www.neotcrdb.com/.

## Introduction

Neoantigens arise from somatic mutation events that are exclusively present in tumor cells and are considered ideal targets for immunotherapy [1]. Neoantigen-based immunotherapies mediate durable responses and long-term remission even in patients with metastatic or advanced cancer [2–5]. These anti-tumor efficacies are considered to be triggered by T cell receptors (TCRs) with specificities for neoantigens presented by human leukocyte antigen (HLA) molecules on tumor cells [6–9]. Therefore, understanding the characteristics of neoantigen-reactive TCRs is critical to interpreting neoantigen-based immune responses and better designing effective immunotherapy for cancer patients.

Recent advances in immunology and high-throughput TCR sequencing approaches have facilitated experimental efforts to map TCR repertoires for a given epitope, including neoantigen [10–12]. The major challenge in this field is the accurate identification of the tumor-reactive, neoantigen-specific TCRs from enormous TCR sequencing data. Although several methods for TCR specificity prediction [13, 14] have been established, these methods do not allow for predicting TCRs specific for neoantigen, largely due to the scarcity of training data computationally connecting a given TCR sequence to a specific neoantigen. Biological validation, such as the *in vitro* assays based on T-cell recognition of targets expressing specified peptide-HLA molecules (peptide-HLA multimers) and specific cytotoxic trails using TCR-engineered T cells, remain the main methods for obtaining neoantigen-specific TCRs. However, this is a time-consuming, difficult task. Moreover, a particular neoantigen may trigger several DNA rearrangement processes, leading to a diversity of TCR repertoires; and a particular neoantigen-specific TCR may resurface in various cancer patients with the same mutation [15, 16]. Additionally, the products submitted for TCR sequencing may contain bystander T cells with viral specificities [9], which makes it more challenging to isolate TCRs targeting neoantigens. To date, several databases of TCR sequences have been developed, such as VDJdb [17, 18], McPAS-TCR [19], and TCRdb [20]. These databases contain a large number of TCR sequences from humans and other species, making them significant resources. VDJdb and McPAS-TCR provide the information of TCR-related antigens as well. However, each TCR sequence in the VDJdb is α or β chain, not paired αβTCR. McPAS-TCR provides paired αβTCR, but the majority of these are associated to viruses. TCRdb contains more than 277 million highly reliable TCR sequences that extracted from TCR-Seq data but without the information of related antigens and HLA alleles. None of these TCR-containing databases pay particular attention to TCR sequences that are specific to neoantigens, especially the TCRs with entire α and β chains that can be directly used in TCR-T cell-based immunotherapy for cancer patients in the clinic. Therefore, the scattered data of functionally verified neoantigen-specific TCRs is urgently required to be gathered in a specialized database and made easy to assess and evaluate.

This study presents NeoTCR, an immunoinformatic database of TCR sequences associated with numerous neoantigens in diverse malignancies, based on published literature. NeoTCR contains comprehensive details on each TCR sequence, such as the related neoantigen and corresponding neoepitope, the restriction of the HLA allele, the tissue of origin, and the reported literature. To quickly identify neoantigen-specific TCRs from raw sequencing data, NeoTCR offers a one-stop analysis and user-friendly solution for annotation and visualization of TCR clonotypes, identification of existing neoantigen-specific TCRs, and exclusion of bystander viral-associated TCRs. NeoTCR is a useful database for examining the activity of neoantigen-associated T-cells in antitumor immunity, which can expand our knowledge of the biological functions and applications of neoantigen-specific TCRs.

## Implementation

The platform of the NeoTCR database was based on the web and B/S architecture, which was implemented using Spring Boot and VUE. All TCR sequences related to neoantigens were stored in MySQL. The user interface was developed using Element UI and the data visualization was developed using ECharts which provide intuitive figures to describe the follow-up analysis result (Figure 1a).

**Figure 1.**
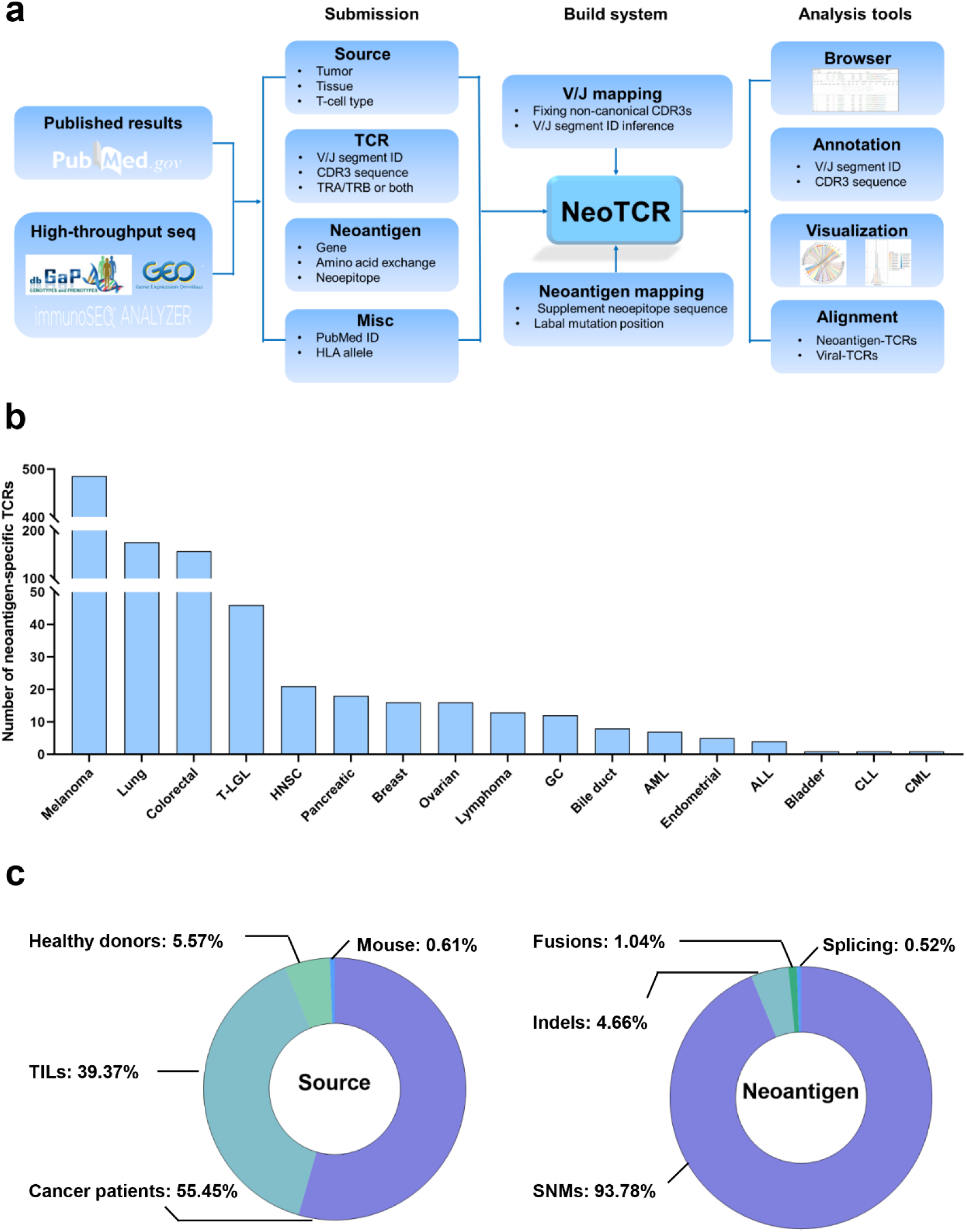
Overview of NeoTCR. **a.** Overall design and construction of NeoTCR. **b.** Data summary and neoantigen-specific TCRs (T cell receptors) across different cancer types. **c.** The source of neoantigen-specific TCRs and TCR-related neoantigen compositions in NeoTCR.

## Database content and usage

### Overview of NeoTCR and data summary

Currently, NeoTCR contains 988 entries, curated from 52 publications, of TCR sequences that were found in eighteen types of cancer, including solid tumors and hematologic malignancies (Figure 1b). As shown in the pie charts of Figure 1c, more than 99% of the sequences are from human data, including peripheral lymphocytes of cancer patients (*N* = 538 entries) or healthy donors (*N* = 55), tumor-infiltrating lymphocytes (TILs, *N* = 389), and the rest come from humanized mouse models (*N* = 6). The TCR-associated neoantigens in NeoTCR contain all types of somatic mutations, most are single nucleotide mutation (SNM, *N* = 181), the others including insertion and deletion (indel, *N* = 9), splicing (*N* = 1), and fusion (*N* = 2).

### Functional description of NeoTCR

To explore the characteristics of neoantigen-specific TCRs, NeoTCR provides three functional modules (Figure 2): (i) Browse section, allowing users to discover and compare interested TCR sequences in different conditions; (ii) Annotation section, enabling one-stop analysis of users’ own TCR sequencing data to extract general features of TCR repertoires, and interactive data visualization of the annotation results, describing the TCR repertoire in TCR diversity, complementarity-determining region 3 (CDR3) length distribution, frequency of CDR3 sequences, usage frequency of V/J gene segment, and V-(D)-J gene segment utilization; (iii) Alignment section, aligning CDR3 sequences with NeoTCR and other existing TCR-databases, typically viral-associated, to identify the known neoantigen-specific TCRs while simultaneously exclude bystander viral-associated TCRs.

**Figure 2.**
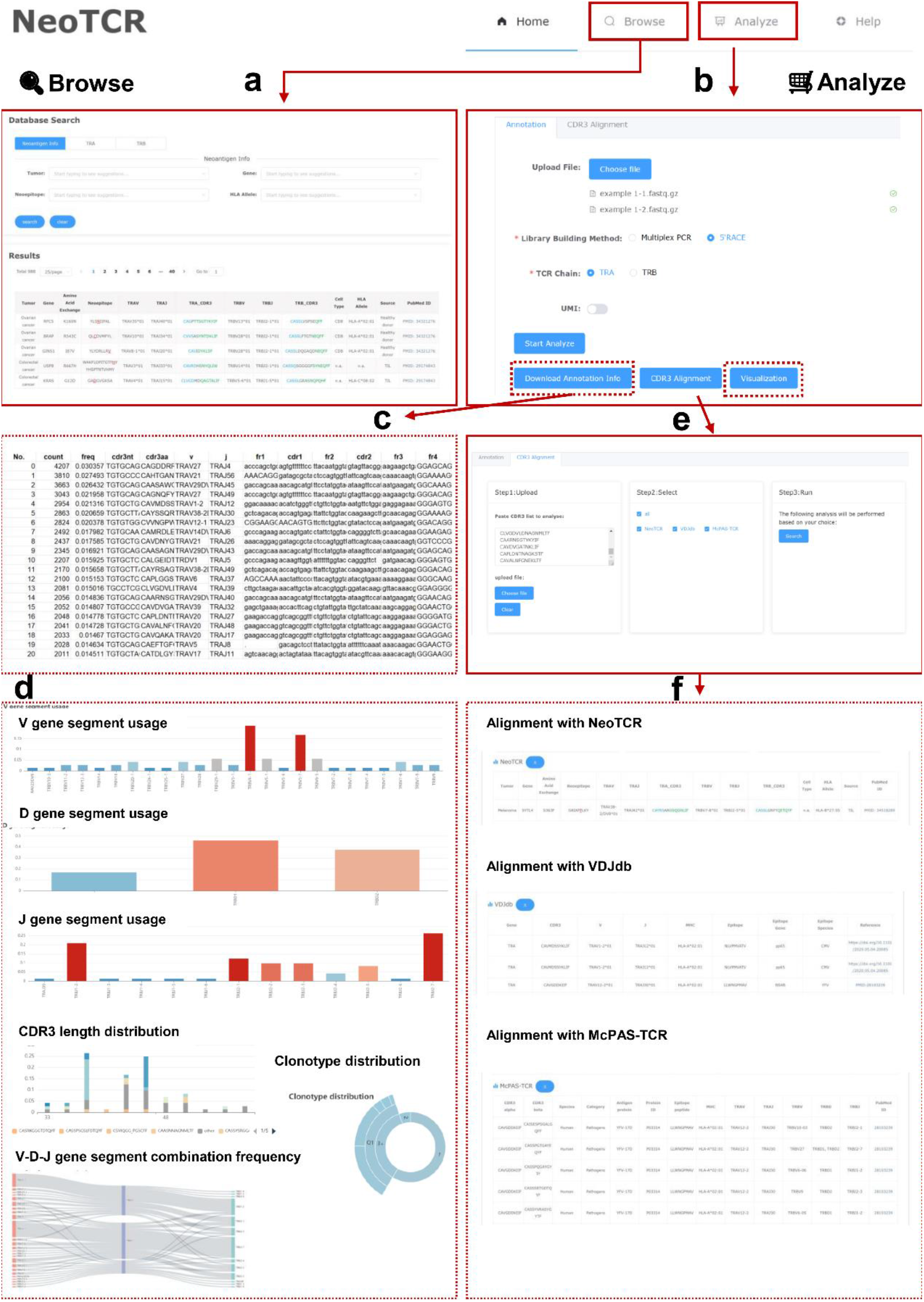
The functional web pages of NeoTCR. **a**. Browse: Users could search their interested TCR sequences and corresponding neoantigens in flexible ways and view the results in user-friendly interactive pages. **b.** Analyze: Users could analyze their own TCR sequencing data in the annotation section and download the annotation information in.txt format (**c**), view the TCR clonotype in the interactive page (**d**), or visit the CDR3 (complementarity-determining region 3) alignment section (**e**) to analyze the annotated CDR3 sequences with NeoTCR and other datasets in more detail (**f**).

### Application of NeoTCR

#### Performing a specific neoantigen-associated TCRs query

NeoTCR provides a tool to investigate the connections between certain neoantigens and the TCRs that recognize them. As an illustration, we looked for the TCR sequences associated with *TP53*-derived neoantigens. We used a fuzzy search performed by entering *‘TP53’* in the ‘Gene’ query box (Figure 3a). The search retrieved 23 entries of TCR sequences that have been identified in TILs from patients with non-small cell lung cancer, ovarian cancer, or colorectal cancer. These TCRs were specific to 11 neoepitopes arising from 6 single nucleotide mutations of *TP53* (R248L/T172I/G245S/R175H/Y220C/R248W), respectively, and thus exhibited different HLA restrictions (Figure 3b). Of striking interest was that different cancer patients with the same HLA allele (HLA-A*68:01) might have potent anti-tumor TCR rearrangement at the same somatic position in R248 but with a different amino acid exchange (L or W). Based on the scant information available, we found that the majority of R248L-specific TCR sequences shared similar TRAV/TRBV gene segments but varied in their TRAJ/TRBJ gene segments. Unlike this, the HLA-A*68:01-restricted, R248W-reactive TCRs differed in their TRAJ and TRBV gene segments while sharing the identical TRAV and TRBJ gene segments (Figure 3C). The V and J gene segments of α or β chains in R175H, however, did not show any similarities (Table 1). For a single mutation of *TP53*, the CDR3 of TRA or TRB were all different, despite the V or J gene segments’ similarity. It was suggested by intriguing differences in the CDR3 sequences that a single mutation may result in a large rearrangement of TCRs.

**Figure 3.**
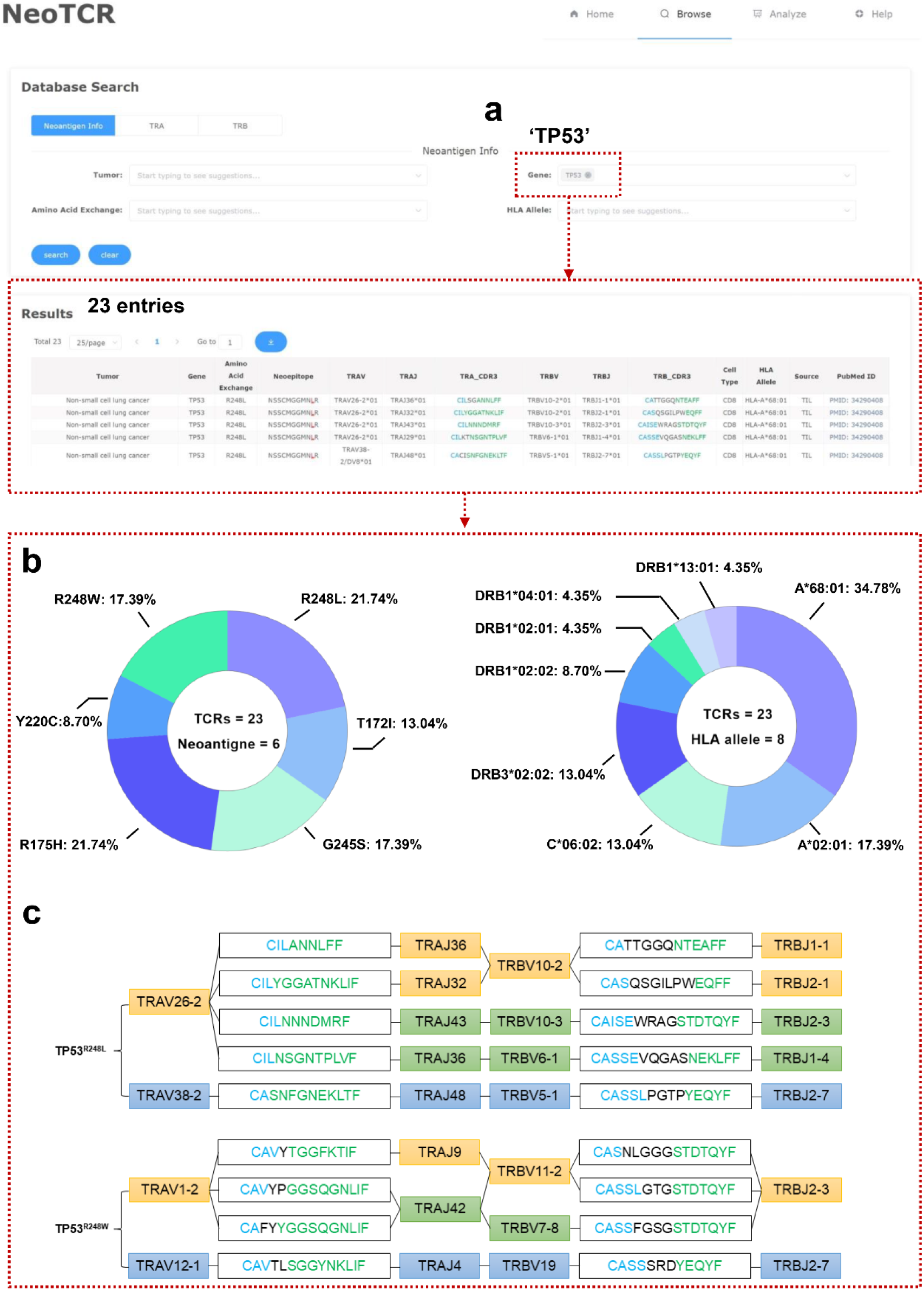
Performing a particular neoantigen-associated TCRs search. **a**. The search for the keyword ‘*TP53*’ returned 23 results. **b.** The distribution of neoantigen and HLA (human leukocyte antigen) allele in *TP53* mutation-specific TCRs. **c.** TCR sequence characteristics that target *TP53*^R248L^ and *TP53*^R248W^.

**Table 1.** Characteristics of TCR sequences specific to *TP53*^R175H^

| Neoepitope | V gene | J gene | CDR3 sequence | HLA allele | PMID |
| --- | --- | --- | --- | --- | --- |
| HMTEVVR<u>H</u>C | TRAV12-3 | TRAJ12 | CAMSGLKEDSSYLIF | A*02:01 | 30714987 |
|  | TRBV27 | TRBJ2-3 | CASSIQQGADTQYF |  |  |
| VVR<u>H</u>CPHHERCSDSD, | TRAV24 | TRAJ10 | CALITGGGNKLTF | DRB1*13:01 | 30714987 |
| QHMTEVVR<u>H</u>CPHHER | TRBV6-2 | TRBJ1-6 | CASRLQGWSPLHF |  |  |
| HMTEVVR<u>H</u>C | TRAV12-1 | TRAJ13 | CVVGGYQKVTF | A*02:01 | 30709841 |
|  | TRBV6-1 | TRBJ2-7 | CASSEEQYF |  |  |
| HMTEVVR<u>H</u>C | TRAV6 | TRAJ43 | CALDDMRF | A*02:01 | 30709841 |
|  | TRBV11-2 | TRBJ2-2 | CASSLTGELFF |  |  |
| HMTEVVR<u>H</u>C | TRAV38-1 | TRAJ28 | CAFMYSGAGSYQLTF | A*02:01 | 30709841 |
|  | TRBV10-3 | TRBJ1-6 | CAISESPLHF |  |  |

#### TCR repertoire annotation and clonotype visualization

To demonstrate the utility of NeoTCR in annotating high-throughput TCR sequencing data, we used bulk-TCRα sequencing data from neoantigen-induced T-cell subsets isolated from peripheral blood mononuclear cells of a cancer patient as an example. Bulk-TCR sequencing data was loaded using the ‘Upload file’ option in the ‘Analyze-Annotation’ window. We then selected the ‘5’ Race’ option and clicked ‘Start Analyze’. The raw sequencing data revealed 202 TCR-α sequences, each representing 0.000072 to 3.036 percent of the total sequences (Table S1). Visualizing results showed that CDR3 length was primarily restricted to 30-45 amino acids (Figure 4a). The top 5 CDR3 sequences were ‘CAGDDRRGGYNKLIF’, ‘CAHTGANSKLTF’, ‘CAASAWGGADGLTF’, ‘CAGNQFYF’, and ‘CAVMDSSYKLIF’ (Figure 4b). The most used V and J gene segments were TRAV29/DV5 and TRAJ45, respectively (Figure 4c and 4d), and these two gene segments had the highest combination frequency as well (Figure 4e).

**Figure 4.**
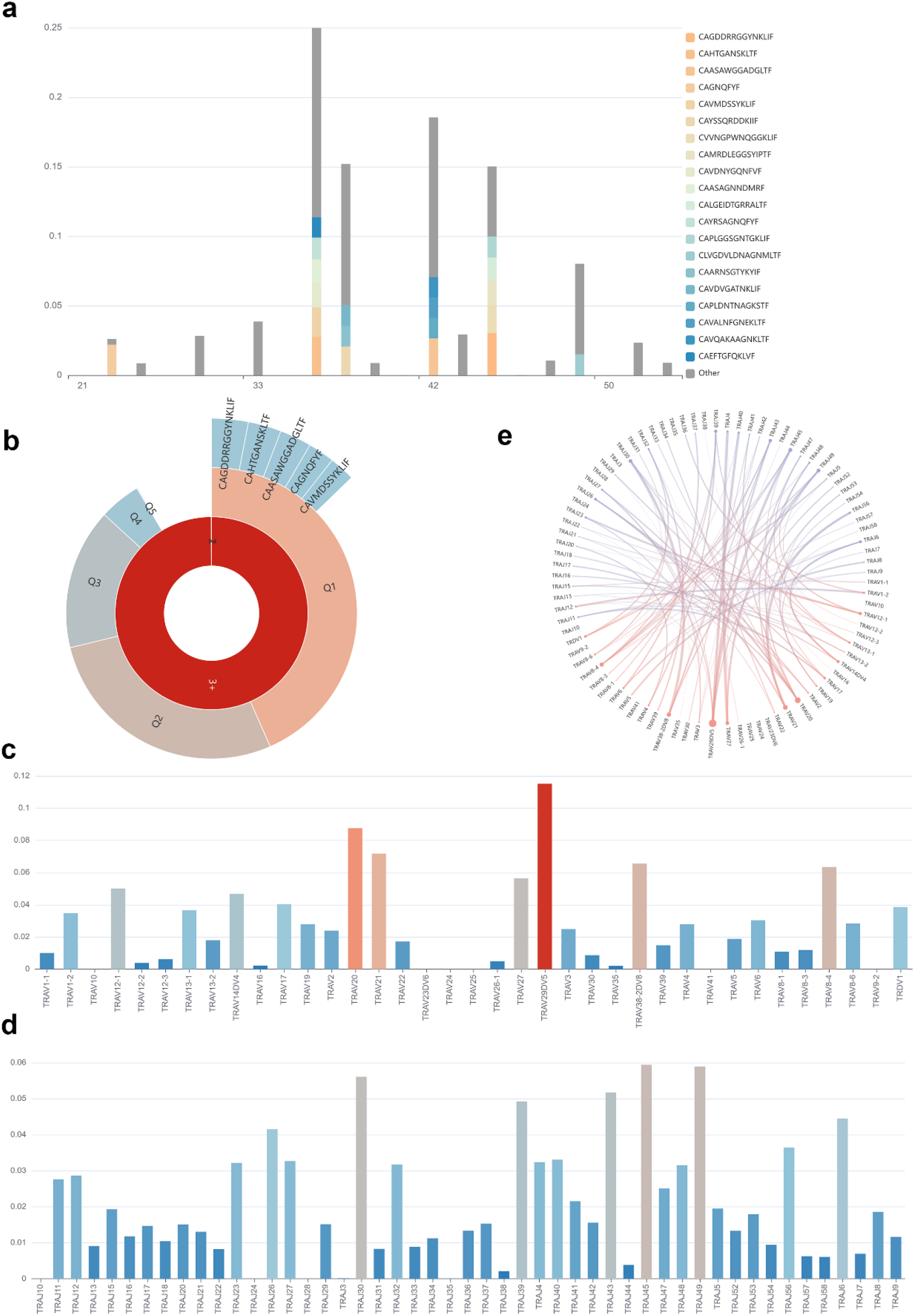
Visualization of the TCR annotation results. **a.** Length distribution of CDR3 sequences. **b**. Frequency of CDR3 sequences. **c.** Usage frequency of V gene segment. **d.** Usage frequency of J gene segment. **e.** Combination possibility of V and J gene segments.

#### CDR3 sequences alignment

One of the main focuses of studies on TCRs is connecting a given TCR to the antigens that it binds and recognizes. Distinguishing whether a TCR is neoantigen-specific or viral-associated might significantly reduce verification efforts. CDR3, a product of the TCR rearrangement, is in charge of antigen recognition, which forms the cornerstone of TCR specificity. The annotated CDR3 sequences we received from the example bulk-TCR sequencing data were further examined using the ‘CDR3 Alignment’ section in NeoTCR (Figure 5a). We learned through NeoTCR that ‘CAYRSARGSQGNLIF’ had been shown to target the neoantigen *SYTL4*^S363F^ in an HLA-B*27:05-dependent manner, and the information on corresponding TCR-β was also exhibited in this database (Figure 5b, Table S2). This *SYTL4*^S363F^-specific TCR had been demonstrated to efficiently induce immune responses against cancer cells expressing both *SYTL4*^S363F^ and HLA-B*27:05 [21]. According to this information, T-cell therapy that targets this paired TCR may be appropriate for this cancer patient. In this example data, bystander viral-associated TCRs were intermingled, including those from the cytomegalovirus (CMV), Epstein-Barr virus (EBV), SARS-CoV-2, yellow fever virus (YFV), and influenza A. CMV-associated TCRs were most commonly found in the alignment results from VDJdb and McPAS-TCR, notably those specific to pp65 (Figure 5c, Table S3-S4). 2 and 15 distinct TCRs were only found, respectively, in McPAS-TCR, and VDJdb (Figure 5c).

**Figure 5.**
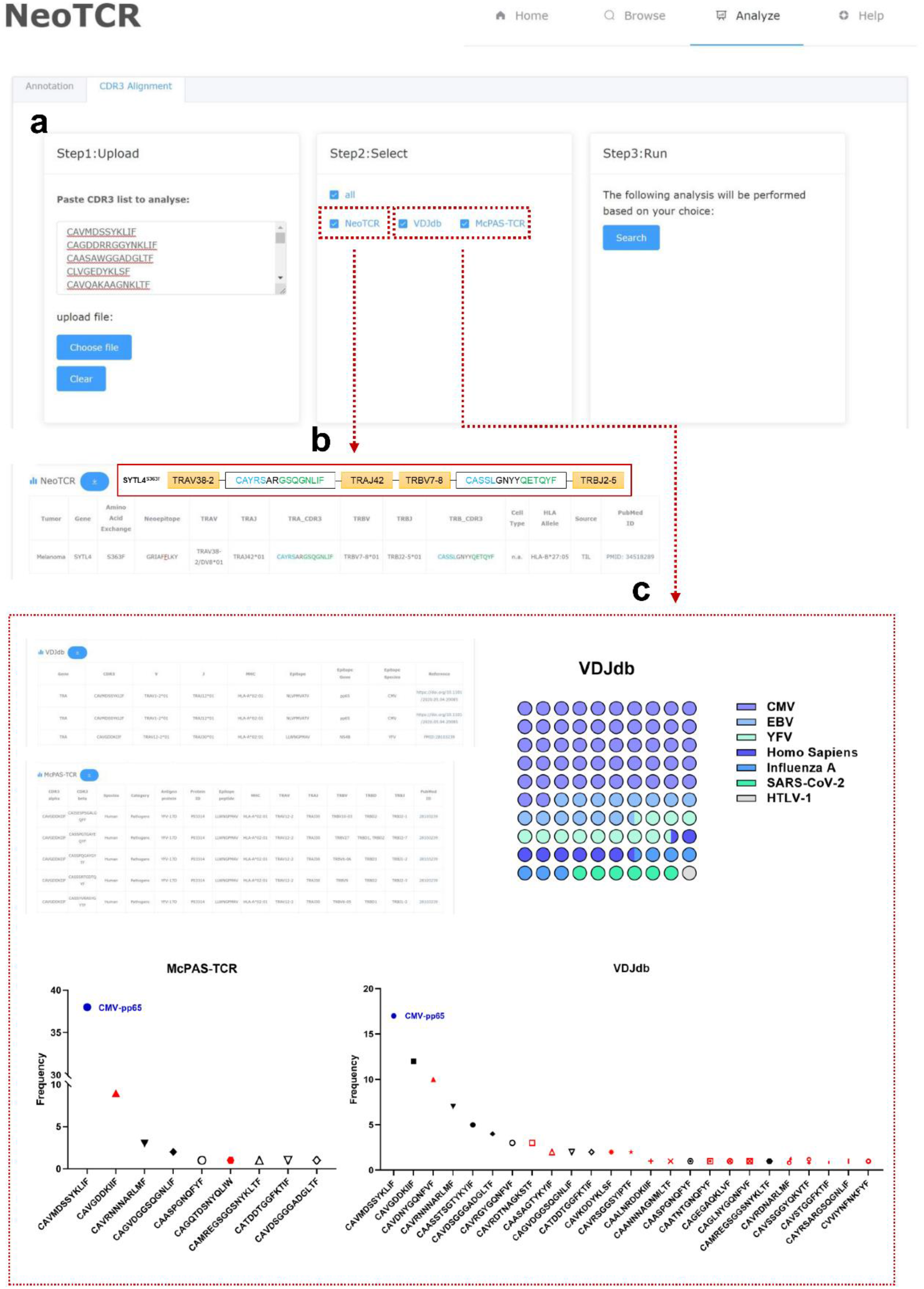
CDR3 sequences alignment. **a.** An illustration of an alignment using the TCRα sequences that were annotated from the sample raw bulk-TCR sequencing data. **b.** A neoantigen-specific TCR sequence had been found in NeoTCR, as well as the corresponding β chain sequence. **c.** Bystander viral-associated TCR sequences were intermingled with different frequencies in this example data.

## Methods

### Data collection and processing

To build NeoTCR, a PubMed query was conducted with variations of the following search terms: ((T cell receptor [Title / Abstract]) OR (TCR [Title / Abstract]) OR (CDR3 sequence [Title / Abstract]) OR (T cell repertoire [Title / Abstract]) OR (T cell [Title / Abstract]) OR (tumor-infiltrating lymphocytes [Title / Abstract]) AND (neoantigen). Further publications were added based on citations or similar articles found using the PubMed search. For the TCR sequences that lack the variable (V) and/or joining (J) specifications, or contain incomplete or excessive CDR3 sequences, the raw data (if available) was downloaded and submitted to the pipeline built in this database to generate the annotated information. Generally, NeoTCR collects neoantigen-related data from published experimental works.

### Data organization

Each entry in the database contains one TCR sequence identified in one study, which is described by the following fields:

#### Tumor

Disease in which the TCR sequence was identified.

#### Gene, Amino Acid Exchange, and Neoepitope

These fields describe the characteristics of TCR-associated neoantigens: Antigen – the antigen protein that the TCR target; Amino Acid Exchange – the amino acid somatic from the corresponding wild-type antigen protein, including SNM, indel, fusion, and splicing; Neoepitope – the amino acid sequence of neoepitope that TCR binds and recognizes, the mutated amino acid is underscored.

#### TRAV, TRAJ, TRA_CDR3 and TRBV, TRBJ, TRB_CDR3

The information of TCRα and TCRβ chain. TRAV, TRBV – The V gene segment ID of TCRα and TCRβ, respectively. TRAJ, TRBJ – The J gene segment ID of TCRα and TCRβ, respectively. Each segment ID is with strictly IMGT nomenclature. TRA_CDR3, TRB_CDR3 – The CDR3 amino acid sequence of TCRα and TCRβ, respectively. The complete CDR3 sequence is starting with the conserved cysteine (C) at the 5’ end of the V segment and ends with the conserved phenylalanine (F) or Tryptophane (W) at the 3’ end of the J segment. Trimmed or excessive sequences will be fixed at the data processing stage in case sufficient V/J germline parts are present.

#### Cell Type

This field describes the lymphocyte subtype of the TCR, e.g. CD4 and CD8.

#### HLA Allele

The HLA restriction of the TCR-associated neoepitope, as described in the original article.

#### Source

This field describes the tissues that the TCR sequence found, e.g. Healthy donor (peripheral lymphocytes of the healthy donor), TIL (tumor-infiltrating lymphocytes), Patient (peripheral lymphocytes of the cancer patient), and humanized mouse models of tumors.

#### PubMed ID

The PubMed Identifier (PMID) of the study’s entry in the PubMed Biomedical Literature Citation Database (https://pubmed.ncbi.nlm.nih.gov/). A link is provided for easy access to the paper.

### Database analysis tools

#### Annotation tool

A uniform pipeline to annotate TCRα and TCRβ chains was developed in NeoTCR as described below. First, raw reads of TCR sequencing were trimmed by Trimmomatic [22] to remove low-quality bases, unknown bases labeling N, and adapters contamination. A quality control test was performed by FastQC [23]. Then UMI-tools [24] was used to process the data containing unique molecular identifiers (UMIs), which enabled the correction of polymerase chain reaction (PCR) amplification biases and quantification of the number of receptors expressed. To generate the annotated information, MiXCR [25] was employed to align sequencing reads to reference V, D, J, and C segments of the TCR chain and assemble clonotypes.

#### Visualization tool

The follow-up analysis of the annotated information was performed using Python and ECharts [26] to implement data visualization. Briefly, the annotated data was grouped into statistics in different dimensions to analyze the TCR repertoires. The visualization, including the counts of V, (D), and J genes of each TCR chain, the V-J genes usage of TCRα chain, the V-D-J genes usage of TCRβ chain, the CDR3 length, and clonotypes distribution was performed through ECharts [26].

#### Alignment tool

To label the known neoantigen-specific TCRs and bystander viral TCRs, the alignment of CDR3 sequences was applied based on several databases of TCRs, including VDJdb [17, 18], McPAS [19], and NeoTCR.

## Discussion

TCR diversity is produced by the random recombination of the V, (D), and J gene segments and the pairing of TCRα and TCRβ chains. There are now a lot of TCRs with neoantigen specificities that were isolated from tumor specimens or peripheral blood [4, 6, 27]. In this regard, we developed the extensive database NeoTCR, which is dedicated to housing the TCR sequences connected to neoantigens.

We anticipate that NeoTCR database and the web server can facilitate biomedical researchers in developing novel ideas on neoantigen-induced immune responses as well as clinicians in designing personalized TCR-T cell-based immunotherapy for cancer patients. For instance, researchers can compare TCR repertoires with the same CDR3 sequences (e.g. paired TCRs have the same TRA_CDR3 sequence but different TRB_CDR3; or the TCRα/β chains have the same CDR3 sequence but different V and/or J gene segments). For a more specific individual condition, if an HLA-DRB3*02:02-positive patient harbors *TP53*^G245S^ mutation, the clinicians can search NeoTCR to quickly find a specific TCR (TRAV8-1*01 – CAVKGDYKLSF – TRAJ20*01; TRBV11-2*01 – CASSLVNTEAFF – TRBJ1-1*01) that satisfies these requirements, which can be employed in TCR-T cell-based immunotherapy for this patient. Besides, NeoTCR offers two web tools for analyzing raw TCR-sequencing data: *Annotation*: Extracting general features of TCR repertoires and further visualizing the clonotypes. *CDR3 Alignment:* Aligning the annotated CDR3 sequences with NeoTCR and other existing TCR databases, typically viral-associated, to find out the known neoantigen-specific TCRs and to exclude bystander viral-associated TCRs.

There will be a fast-rising number of pertinent research and more neoantigen-related TCR sequences will be discovered as a result of the increased interest in the involvement of neoantigens in TCR rearrangement and cancer immunotherapies. To expand the database’s storage capacity, we are working to acquire more neoantigen-specific TCR sequences from raw high-throughput sequencing data and periodically update NeoTCR with newly discovered TCRs. Additionally, we intend to build a model to forecast neoantigen-TCR binding for human TCRs based on deep learning techniques trained on neoantigen-specific TCR data gathered from NeoTCR and other related datasets.

## Data availability

NeoTCR is a comprehensive online database available at http://neotcrdb.com.

The guidelines for NeoTCR are available at http://www.neotcrdb.com/help.

## CRediT author statement

**Weijun Zhou**: Conceptualization, Validation, Investigation, Resources, Data Curation, Writing-Original Draft, Writing-Review & Edition, Visualization, Project administration, Funding acquisition. **Wenting Xiang**: Methodology, Software, Data Curation, Writing-original Draft, Writing-Review & Edition. **Jinyi Yu**: Resources. **Yichen Pan**: Data Curation. **Zhihan Ruan**: Software. **Kankan Wang**: Writing-Original Draft, Writing-Review & Edition, Visualization, Project administration, Funding acquisition. **Jian Liu**: Conceptualization, Writing-Original Draft, Writing-Review & Edition, Visualization, Project administration.

## Acknowledgments

This work was partially supported by the National Natural Science Foundation of China (grant number 81890994), the National Key R&D Program of China [grant number 2019YFA0905902]; the Guangdong Basic and Applied Basic Research Foundation, China (grant number 2019A1515010299), the Science and Technology Program of Guangzhou, China (grant number 202102020727), the Innovative Research Team of High-level Local Universities in Shanghai, and the Dean’ fund of Zhujiang Hospital of Southern Medical University, China (grant number yzjj2019qn05). The authors are grateful to Kellie N Smith for the helpful sharing of TCR sequencing data. We thank the developers of VDJdb and McPAS-TCR for the continuous development of their respective databases over the past several years.

## Supplementary Information

**Table S1.**

Annotation results of the example TCR sequencing data.

**Table S2 - S4.**

The CDR3 sequences alignment results from NeoTCR (Table S2), VDJdb (Table S3), and McPAC-TCR (Table S4).

## References

1. Richters MM, Xia H, Campbell KM, Gillanders WE, Griffith OL, Griffith M. Best practices for bioinformatic characterization of neoantigens for clinical utility. Genome Med 2019;11:56.

2. Carlino MS, Larkin J, Long GV. Immune checkpoint inhibitors in melanoma. Lancet 2021;398:1002–1014.

3. Tran E, Robbins PF, Lu YC, Prickett TD, Gartner JJ, Jia L, et al. T-Cell transfer therapy targeting mutant KRAS in cancer. N Engl J Med 2016;375:2255–2262.

4. Tran E, Turcotte S, Gros A, Robbins PF, Lu YC, Dudley ME, et al. Cancer immunotherapy based on mutation-specific CD4+ T cells in a patient with epithelial cancer. Science 2014;344:641–645.

5. Zacharakis N, Chinnasamy H, Black M, Xu H, Lu YC, Zheng Z, et al. Immune recognition of somatic mutations leading to complete durable regression in metastatic breast cancer. Nat Med 2018;24:724–730.

6. Caushi JX, Zhang J, Ji Z, Vaghasia A, Zhang B, Hsiue EH, et al. Transcriptional programs of neoantigen-specific TIL in anti-PD-1-treated lung cancers. Nature 2021;596:126–132.

7. Krishna S, Lowery FJ, Copeland AR, Bahadiroglu E, Mukherjee R, Jia L, et al. Stem-like CD8 T cells mediate response of adoptive cell immunotherapy against human cancer. Science 2020;370:1328–1334.

8. Kristensen NP, Heeke C, Tvingsholm SA, Borch A, Draghi A, Crowther MD, et al. Neoantigen-reactive CD8+ T cells affect clinical outcome of adoptive cell therapy with tumor-infiltrating lymphocytes in melanoma. J Clin Invest 2022;132:e150535

9. Lowery FJ, Krishna S, Yossef R, Parikh NB, Chatani PD, Zacharakis N, et al. Molecular signatures of antitumor neoantigen-reactive T cells from metastatic human cancers. Science 2022;375:877–884.

10. Calis JJ, Rosenberg BR. Characterizing immune repertoires by high throughput sequencing: Strategies and applications. Trends Immunol 2014;35:581–590.

11. Friedensohn S, Khan TA, Reddy ST. Advanced methodologies in High-Throughput sequencing of immune repertoires. Trends Biotechnol 2017;35:203–214.

12. Pai JA, Satpathy AT. High-throughput and single-cell T cell receptor sequencing technologies. Nat Methods 2021;18:881–892.

13. Gielis S, Moris P, Bittremieux W, De Neuter N, Ogunjimi B, Laukens K, et al. Detection of enriched t cell epitope specificity in full t cell receptor sequence repertoires. Front Immunol 2019;10:2820.

14. Montemurro A, Schuster V, Povlsen HR, Bentzen AK, Jurtz V, Chronister WD, et al. NetTCR-2.0 enables accurate prediction of TCR-peptide binding by using paired TCRalpha and beta sequence data. Commun Biol 2021;4:1060.

15. Levin N, Paria BC, Vale NR, Yossef R, Lowery FJ, Parkhurst MR, et al. Identification and validation of t-cell receptors targeting RAS hotspot mutations in human cancers for use in cell-based immunotherapy. Clin Cancer Res 2021;27:5084–5095.

16. Yossef R, Tran E, Deniger DC, Gros A, Pasetto A, Parkhurst MR, et al. Enhanced detection of neoantigen-reactive T cells targeting unique and shared oncogenes for personalized cancer immunotherapy. JCI Insight 2018;3:e122467.

17. Shugay M, Bagaev DV, Zvyagin IV, Vroomans RM, Crawford JC, Dolton G, et al. VDJdb: A curated database of T-cell receptor sequences with known antigen specificity. Nucleic Acids Res 2018;46:419–427.

18. Bagaev DV, Vroomans R, Samir J, Stervbo U, Rius C, Dolton G, et al. VDJdb in 2019: Database extension, new analysis infrastructure and a T-cell receptor motif compendium. Nucleic Acids Res 2020;48:1057–1062.

19. Tickotsky N, Sagiv T, Prilusky J, Shifrut E, Friedman N. McPAS-TCR: A manually curated catalogue of pathology-associated T cell receptor sequences. Bioinformatics 2017;33:2924–2929.

20. Chen SY, Yue T, Lei Q, Guo AY. TCRdb: A comprehensive database for T-cell receptor sequences with powerful search function. Nucleic Acids Res 2021;49:468–474.

21. Braunlein E., Lupoli G., Fuchsl F., Abualrous E. T., de Andrade Kratzig N., Gosmann D., Wietbrock L., Lange S., Engleitner T., Lan H., Audehm S., Effenberger M., Boxberg M., Steiger K., Chang Y., Yu K., Atay C., Bassermann F., Weichert W., Busch D. H., Rad R., Freund C., Antes I., Krackhardt A. M. Functional analysis of peripheral and intratumoral neoantigen-specific TCRs identified in a patient with melanoma. J Immunother Cancer. 2021. 9: e002754.

22. Bolger AM, Lohse M, Usadel B. Trimmomatic: A flexible trimmer for Illumina sequence data. Bioinformatics 2014;30:2114–20.

23. de Sena BG, Smith AD. Falco: High-speed FastQC emulation for quality control of sequencing data. F1000Res 2019;8:1874.

24. Smith T, Heger A, Sudbery I. UMI-tools: Modeling sequencing errors in Unique Molecular Identifiers to improve quantification accuracy. Genome Res 2017;27:491–499.

25. Bolotin DA, Poslavsky S, Mitrophanov I, Shugay M, Mamedov IZ, Putintseva EV, et al. MiXCR: Software for comprehensive adaptive immunity profiling. Nat Methods 2015;12:380–381.

26. Deqing L, Honghui M, Yi S, Shuang S, Wenli Z, Junting W, et al. ECharts: A declarative framework for rapid construction of web-based visualization. Visual Informatics 2018;2:136–146.

27. Yossef R, Tran E, Deniger DC, Gros A, Pasetto A, Parkhurst MR, et al. Enhanced detection of neoantigen-reactive T cells targeting unique and shared oncogenes for personalized cancer immunotherapy. JCI Insight 2018;3:e122467.

